# Distinct neural mechanisms and temporal constraints govern a cascade of audiotactile interactions

**DOI:** 10.1101/446112

**Authors:** Johanna M. Zumer, Thomas P. White, Uta Noppeney

## Abstract

Asynchrony is a critical cue informing the brain whether sensory signals are caused by a common source and should be integrated or segregated. It is unclear how the brain binds audiotactile signals into behavioural benefits depending on their asynchrony. Participants actively responded (psychophysics) or passively attended (electroencephalogrpahy) to noise bursts, ‘taps-to-the-face’, and their audiotactile (AT) combinations at seven audiotactile asynchronies: 0, ±20, ±70, and ±500ms. Observers were faster at detecting AT than unisensory stimuli, maximally for synchronous stimulation and declining within a ≤70ms temporal integration window. We observed AT interactions for (1) near-synchronous stimuli within a ≤20ms temporal integration window for evoked response potentials (ERPs) at 110ms and ∼400ms, (2) specifically ±70ms asynchronies, across the P200 ERP and theta-band inter-trial coherence (ITC) and power at ∼200ms, with a frontocentral topography, and (3) beta-band power across several asynchronies. Our results suggest that early AT interactions for ERP and theta-band ITC and power mediate behavioural response facilitation within a ≤70ms temporal integration window, but beta-band power reflects AT interactions that are less relevant for behaviour. This diversity of temporal profiles and constraints demonstrates how audiotactile integration unfolds in a cascade of interactions to generate behavioural benefits.

## Introduction

Imagine sitting outside on a summer evening. Suddenly you hear a buzz and then feel a prick to your skin, as the mosquito lands. You are faster to swat it away because you first heard it coming. This faster detection of a multisensory event, known as the redundant target effect (RTE) (Miller 1982; Diederich and Colonius 2004; Sperdin et al. 2009), illustrates the enormous benefits of multisensory integration.

Importantly, we should integrate signals only if they arise from a common source but segregate them otherwise. Synchrony is a critical cue for indicating whether two signals come from a common source. Multisensory signals need to co-occur within a certain tolerance of asynchrony, termed a temporal integration window (TIW) (Diederich and Colonius 2004). In particular, the RTE typically follows an inverted U-shape function (Blurton et al. 2015) that is maximal for (near)-synchronous signals and tapers off with increasing asynchrony thereby moulding the TIW. Likewise, observers’ perceived synchrony, the emergence of cross-modal biases, and perceptual illusions follow a similar inverted U-shape function with its exact shape varying across different behavioural measures and task-contexts (van Wassenhove et al. 2007; Megevand et al. 2013; Berger and Ehrsson 2014; Donohue et al. 2015).

At the neural level, multisensory influences have been identified in terms of response enhancements and suppressions, super-additive and sub-additive interactions (Meredith and Stein 1983; Stanford et al. 2005; Werner and Noppeney 2010b), shortened neural response latencies (Rowland and Stein 2007) and altered neural representations (Fetsch et al. 2011; Rohe and Noppeney 2015; 2016; Aller and Noppeney 2019; Rohe et al. 2019). Evidence from neuroimaging, neurophysiology, and neuroanatomy has shown that multisensory influences emerge at early and late stages of neural processing (Foxe et al. 2000; Lutkenhoner et al. 2002; Murray et al. 2005; Senkowski et al. 2008; Sperdin et al. 2009; Stekelenburg and Vroomen 2009; Mercier et al. 2013; Mercier et al. 2015) nearly ubiquitously in neocortex (Schroeder and Foxe 2002; Ghazanfar and Schroeder 2006; Lakatos et al. 2007; Werner and Noppeney 2010a; Ibrahim et al. 2016; Atilgan et al. 2018). They arise already at the primary cortical level and increase progressively across the sensory processing hierarchy (Foxe and Schroeder 2005; Bizley et al. 2007; Kayser et al. 2007; Dahl et al. 2009). This multi-stage and multi-site account of multisensory interplay raises the question of whether the myriad multisensory influences is governed by similar neural mechanisms and temporal constraints. Further, how do those neural effects relate to the TIW defined by behavioural indices? Given previous unisensory research showing an increase in the TIW along the sensory processing hierarchy (Hasson et al. 2008; Kiebel et al. 2008), one may for instance hypothesise that early multisensory interactions are confined to narrower temporal integration windows than those occurring at later stages in higher order association cortices (Werner and Noppeney 2011). Moreover, recent neurophysiological studies suggest that multisensory interactions depend on the phase of ongoing neural oscillations and/or rely on mechanisms of phase resetting. For instance, Lakatos et al. (2007) showed that a tactile signal can reset the phase of ongoing oscillations in auditory cortices, but only for specific asynchronies.

The current study aims to define the temporal constraints of multisensory interactions that can be observed for evoked response potentials (ERP), inter-trial coherence (ITC), and induced power responses and relate those to the TIW derived from behavioural response facilitation. Participants were presented with brief airpuff noise bursts, ‘taps to the face’, and their audiotactile (AT) combinations at seven levels of asynchrony: 0, ±20, ±70, and ±500 ms. In the psychophysics study observers were instructed to respond to all A, T, and AT events in a redundant target paradigm; in the EEG study a passive stimulation design was used to avoid response confounds. We then compared the multisensory influences in terms of multisensory interactions (i.e. AT + No stimulation ≠ A + T) across AT asynchrony levels for ERPs, ITC, and time-frequency (TF) power responses and characterised their topography across post-stimulus time.

## Materials and Methods

### Participants

Twenty-five healthy, adult participants with no neurological disorder were recruited from the local university population (students as well as members of the general public) (N=25, 12 female and 13 male; aged between 18-35 years old). One participant was excluded due to an abnormal finding in the structural MRI. Two participants were excluded from the behavioural analysis, because data were not collected for all conditions. Two different participants were excluded from the EEG analysis, because insufficient EEG data were collected. As a result, we included 22 participants in both the behavioural and EEG analysis. They gave written informed consent and were compensated with either cash or course credit. Ethical approval for the study was given by the University of Birmingham Science, Technology, Engineering, and Mathematics Review Committee with approval number ERN_11-0429AP22B.

### Stimulation

Tactile stimulation consisted of a touch to the left side of the face with 200 ms duration. Tactile stimulation to the face was used as an ecologically valid stimulus that requires a rapid response in everyday life. We also chose stimulation to the face (in contrast to hands), as this body location does not require additional processing (e.g. reference frame transformations across the senses) of being potentially crossed relative to body position, thus potentially amenable to a quicker and more automatic route. The auditory association areas that receive feed-forward (layer 4) input from somatosensory stimulation appear to be optimally stimulated by cutaneous stimulation of the head and neck (Fu et al. 2003). The left side was chosen based on previous findings that MSI is enhanced with left-side stimulation and right hemisphere involvement (Giard and Peronnet 1999; Downar et al. 2000; Molholm et al. 2002; Hoefer et al. 2013). The part of the face touched was on/near the border between the maxillary (V2) and mandibular (V3) divisions of the trigeminal cranial nerve. A fibre optic cable (part of a fibre optic system: Keyence series FS-N, Neu-Isenburg, Germany) was attached to a Lego pneumatic cylinder and driven to move by pressurised air. The tip of this cable (3 mm diameter) was positioned near the face using a flexible plastic snap-together ‘goose-neck’ pipe that was attached to an adjustable stand. The air pressure changes were controlled by a microcontroller connected via USB to the stimulus computer; communication to the microcontroller was sent via serial port commands in MATLAB (Mathworks, Inc.). The duration of the open valve (i.e. when the diode was extended forward to touch the skin) was set to 200 ms, an ecologically/environmentally valid duration. The fibre optic cable contained a dual fibre: one fibre projected light and the other was a photodiode that detected the light reflectance; from this, the reflectance dynamics confirmed the exact timing of the touch to the skin. As this was a mechanical air-pressure-driven device, it does not have an immediate on/off time (see Figure S1 for a plot of the reflectance data for one trial). This tactile apparreatus was very similar to that used by Leonardelli et al. (2015). After the experiment, subjects were queried as to whether they could hear any noises of the tactile device and none reported that they could.

The auditory stimulus (target) was an airpuff noise of 200 ms duration. The volume of the target was well above threshold for detection but not painfully loud; the volume was stronger on the left channel than on the right (interaural intensity difference) to create the perception of coming from the left. A constant background noise of a recording of a magnetic resonance imaging (MRI) echo-planar imaging sequence (obtained from http://cubricmri.blogspot.co.uk/2012/08/scanner-sounds.html) was played to help mask external noises including those made by the tactile stimulator and for comparison with potential future functional MRI studies. The volume of the background noise, equally loud in both ears, was played at a level comfortable to participants and such that the tactile noises could not be heard. All sounds were presented via E-A-RTone earphone (10 Ohm; E-A-R Auditory Systems) with plastic tube connection (length = 75 cm) to foam ear insert (E-A-RLink size 3A), which also acted as an earplug against external sounds.

### Experimental design

Participants took part in one psychophysics and one EEG session on separate days (typically 4-6 days gap). The experimental design and stimuli were identical across the two sessions. In the psychophysics session participants responded to the first stimulus in a trial irrespective of sensory modality, as fast as possible via a single key board button (i.e. redundant target paradigm). In the EEG session, participants passively perceived the stimuli without an explicit response in order to examine automatic AT interactions (including during unattentive and drowsy states but excluding sleep stages), in order to avoid motor confounds, and to allow for comparison with sleep, non-responsive patients, etc. The EEG session was acquired one hour before the participant’s usual bedtime as part of a longer sleep study (n.b. sleep data will be reported in a separate communication). To ensure that participants were not asleep, we applied sleep staging and excluded data in actual sleep stages (details below). Hence, in this communication, we focus on multisensory interactions in a low vigilance state that have rarely been studied or reported. Yet, multisensory interactions may be most relevant in low vigilance states to attract observers’ attention to salient events in their environment.

In each session, participants were presented with the following ten trial types: no stimulus (or null) condition (N), tactile alone (T), auditory alone (A), and seven audiotactile (AT) conditions varying in asynchrony (−500 ms, −70 ms, −20 ms, 0 ms, 20 ms, 70 ms, 500 ms) where a ‘negative’ asynchrony refers to A-leading-T (Figure 1a). The audiotactile conditions are referred to by the following abbreviations: AT500, AT70, AT20, AT0, TA20, TA70, TA500, respectively. These asynchronies were chosen to fall either within the behaviourally-defined temporal integration window (TIW) (≤70 ms) based on previous studies (e.g. (Navarra et al. 2007; Harrar and Harris 2008; Nishi et al. 2014)) or outside the TIW (± 500 ms). Hence, this study focused on a coarse characterisation of the temporal integration window across both A-leading and T-leading asynchronies rather than a fine-grained analysis of asynchronies within a small range (e.g. as in (Naue et al. 2011)). Seven AT, unisensory A, T and null-trials were presented, interleaved randomly with an inter-trial interval uniformly distributed between 2.0 – 3.5 s, including both unisensory and audiotactile conditions with varying asynchronies between the sensory stimuli. Each trial type was presented 100 times in each session. Trials were presented in blocks of 250 trials (roughly 11.75 minutes) over four blocks separated by short breaks. In the EEG session we occasionally shortened the blocks, but still presented 1000 trials in total. In the psychophysics session the AT500 and TA500 conditions were not collected for two participants; thus for behavioural results, only the data from the remaining twenty-two participants are included (after exclusion also of one participant for the afore-mentioned structural MRI abnormality).

**Figure 1.**
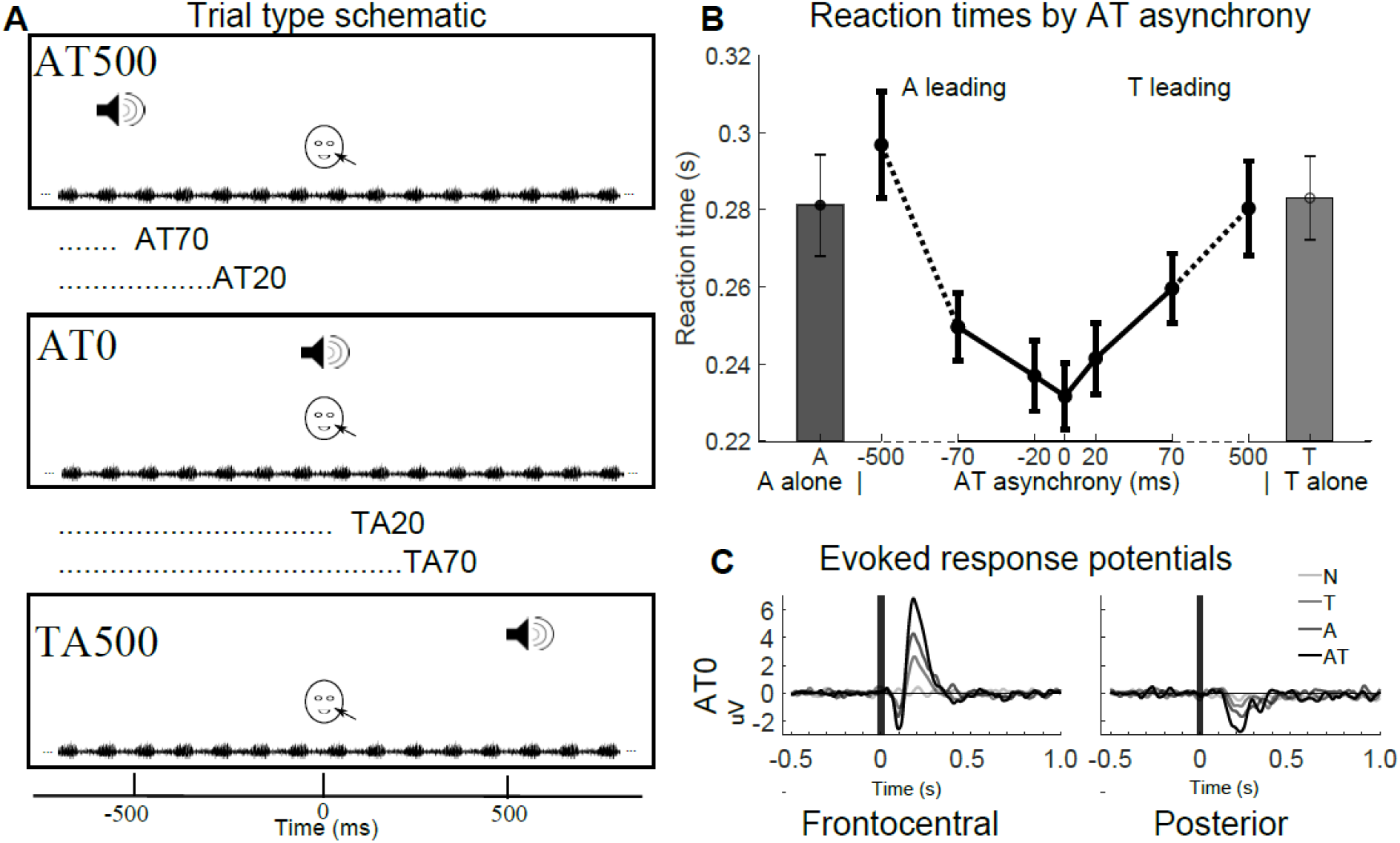
Experimental design, behavioural results, and evoked responses. ***A,*** Each row depicts the onsets of the auditory stimulation (indicated by loudspeaker) and tactile stimulation (indicated by face) for three examples out of the seven AT asynchrony conditions. The wavy line at the bottom indicates the continuous MRI background noise. ***B,*** Reaction times (across subjects’ mean ± SEM). The black lines indicate the AT conditions as a function of AT asynchrony with negative asynchronies indicating auditory-leading; the dark grey and light grey bars indicate the A and T conditions, respectively. ***C,*** Evoked response potentials for N, A, T, and AT0 conditions for frontocentral [‘Fz’ ‘Cz’ ‘F1’ ‘F2’ ‘FC1’ ‘FC2’ ‘C1’ ‘C2’] and posterior [’CP5’ ‘POz’ ‘Pz’ ‘P3’ ‘P4’ ‘C4’ ‘O1’ ‘O2’ ‘P7’ ‘PO7’] sets of sensors. For ERPs for all seven asynchronies in colour, see Figure S3.

Participants kept their eyes closed to obliterate any visual input throughout the experiment. They were seated comfortably with their head stabilised in an adjustable chin rest and were requested to hold their head as still as possible (to promote spatial and temporal consistency of the tactile stimulation over trials).

### EEG recording

EEG data were recorded with a 64 channel BrainProducts MR-compatible cap at 1000 Hz sampling rate, with 63 of the electrodes on the scalp. For all but the first three participants, two additional bipolar electrodes were placed on the face to record horizontal EOG and vertical EOG. For 17 participants, the 64^th^ cap electrode was placed on the participants’ back for recording ECG. For the other 8 participants, the 64^th^ electrode was instead placed on the right (unstimulated) cheek for assistance as EOG/EMG. Signals were digitised at 5000 Hz with an anti-aliasing filter of 1000 Hz, then down-sampled to 1000 Hz with a high-pass filter of 0.1 Hz and low-pass filter of 250 Hz. Electrode impedances were kept below 25 kOhm. Triggers from the stimulus-control computer were sent via LabJack to the EEG acquisition computer.

### Tactile stimulation output

The time course of light reflectance (one example trial depicted in Figure S1) was assessed for each tactile trial to ensure that i. the tactile device actually touched the skin and ii. to determine the touch onset time (1000 Hz sampling rate). After computing the actual onset of the touch from the light reflectance data, subsequently the exact multisensory onset asynchrony was computed for all multisensory trials. Those that deviated by more than ± 5 ms from the desired asynchrony were discarded. This resulted in 16.8% (± 1.1%) and 16.4% (± 1.2%) of trials rejected for the behavioural and EEG data, respectively (N=24, after excluding the participant with structural MRI abnormality).

### Behavioural analysis

After exclusion of trials where touch was not applied or outside the desired asynchrony, sensory trials were additionally discarded with no response or with a response time (RT) faster than 100 ms or slower than 1 s (occurring in total for an average of 2.7±1.1% of trials across conditions). The median RT within a condition for each participant was computed.

For each participant the *redundant target effect* (Hershenson 1962) was computed for each participant by subtracting the median RT of the AT condition at a particular level of asynchrony from the median RT of the fastest (A or T) unisensory condition with the onset of each unisensory condition adjusted for the particular asynchrony (e.g. RT_AT20_ – min(RTT + 20 ms, RT_A_). First, we investigated whether the redundant target effect was different across AT asynchrony conditions, using a one-way repeated-measures ANOVA (rmANOVA) over the seven AT conditions followed by planned post-hoc rmANOVA tests to further narrow the possible asynchrony conditions that drive the overall main effect of asynchrony (i.e. conditions ≤ 70 ms, and if that is significant, then ≤20 ms). Second, we assessed whether the redundant target effect for each condition differed significantly from zero across participants, using a one-sample two-sided t-test.

### EEG analysis: sleep staging

To ensure that only EEG data was used in which participants were awake, given the passive stimulation design with eyes closed and the evening acquisition, standard sleep scoring was performed using American Academy of Sleep medicine (AASM) 2007 criteria in the FASST open-source software (http://www.montefiore.ulg.ac.be/~phillips/FASST.html) (Leclercq et al. 2011) and custom code in MATLAB. Data were segmented into 30 s chunks and referenced to linked-mastoids. Sleep stages were assessed by two of the authors (J.M.Z. and T.P.W.) independently with a correspondence of 88%. Differences were discusssed and a consensus reached (with correspondence of the consensus to each assessor’s scores at 93% and 94%). Any 30 s chunk that was not scored as ‘awake’ was excluded from further analysis. If an individual participant had fewer than 55 trials per condition remaining in the awake stage (prior to artefact rejection), the participant was fully excluded. Two participants were excluded for this reason.

### EEG analysis: preprocessing

All subsequent EEG data processing (after sleep staging) was performed using the open-source toolbox FieldTrip (Oostenveld et al. 2011) (www.fieldtriptoolbox.org) and custom code in MATLAB. Eye movement artefacts were automatically detected using three re-referenced bipolar pairs (‘F7-F8’, ‘Fp2-FT9’, and ‘Fp1-FT10’) and the VEOG if available. These channels’ data were band-pass filtered (1-16 Hz; Butterworth, order 3) and transformed to z-values. The exclusion threshold was set at a z-value of 6 and trials containing these artefacts were excluded. EEG data were re-referenced to the average reference, high-pass filtered (0.2 Hz), band-stop filtered around the line noise and its harmonics (49-51 Hz, 99-101 Hz, and 149-151 Hz), and epoched for each trial. Trials were locked to the onset of the tactile stimulus for tactile and all multisensory conditions and to the auditory or null trigger for A and N conditions, respectively. Initially, the epoch length was from −1.5 s to 2.3 s. Then A trials were shifted ± 0.5, 0.07, 0.02, or 0 s before being added to a T trial, to create the appropriate A+T combination to contrast with AT trials, hence resulting in variable lengths of pre-stimulus and post-stimulus window lengths, depending on the AT asynchrony.

### EEG analysis: multisensory interaction

Multisensory integration in the EEG data was identified in terms of “audiotactile interaction”, i.e. the sum of unisensory (A+T) contrasted to the audiotactile plus null (AT+N). The sum of unisensory (A+T) trials was computed for each AT asynchrony level such that the onsets of the auditory and tactile stimuli were exactly aligned to the trials of the AT condition (i.e. we also accounted for the jitter of tactile onsets, see above). Trials from each condition were randomly sub-selected to ensure an equal number of trials per each of the four conditions in a given contrast (A, T, AT, and N). It is critical to add the null condition (to the multisensory) to account for non-specific effects in a trial such as expectancy of stimulation as well as random noise. Note that removing a pre-stimulus baseline (or, in the present analysis, 0.2 Hz high-pass filter) for removing drift and DC offset is not sufficient to account for these non-specific effects or ‘spontaneous activity’. The argument for a null condition is exemplified in Teder-Salejarvi et al. (2002) and further supported and utilised in other multisensory experiments (Talsma and Woldorff 2005; Bonath et al. 2007; Mishra et al. 2007). Further, the multisensory interaction contrast (with the null-condition included) is equivalent to subtracting the Null condition from each of the stimulus conditions (i.e. an explicit baseline correction): (AT-N)-[(A-N)+(T-N)] = (AT+N)-(A+T). In addition to controlling for the pre-stimulus ‘stimulus expectancy’ confound (as explained in Teder-Salejarvi et al. (2002)), including the Null condition in the interaction contrast also ensures that ‘random noise’ is averaged out similarly for the sum of the two unisensory and the multisensory + Null sum. We also alleviated ‘anticipatory’ or ‘omission’ waves in our data by using a jittered inter-trial interval (uniformly between 2.0-3.5 s).

### EEG analysis: multisensory effects on ERP, inter-trial coherence, and time-frequency power

For the evoked response potential (ERP) analysis, EEG data were low-pass filtered (40 Hz). The average over trials within a participant was computed for the combination of conditions A+T and AT+N separately. We assessed the AT interaction within a 500 ms time window, beginning at the onset of the second stimulus.

For time-frequency analysis, EEG data were Fourier transformed with separate parameters for lower (4-30 Hz) and higher (30-80 Hz) frequencies, with zero padding to a 4 s length (applied to both lower and higher frequencies). Sliding time windows of length equal to four cycles (low frequencies) or 200 ms (high frequencies) at a given frequency in steps of 2 Hz (low frequencies) or 5 Hz (high frequencies), after application of a Hanning taper (low frequencies) or multitaper with +/− 7 Hz smoothing (high frequencies). The complex values were kept for separate analysis of the inter-trial coherence (ITC) (also referred to as phase-locking factor or phase-consistency index) and the time-frequency (TF) power magnitude. Note that the sum of trials of different condition types (i.e. A+T and AT+N) was computed prior to Fourier transformation so that any cancellation due to phase differences would occur prior to obtaining the Fourier complex value (see Senkowski et al. (2007)). The ITC was computed for each condition and subject as the absolute value of the sum of the complex values over trials. We assessed the AT interactions for ITC and TF power separately for ‘low frequency’ and ‘high frequency’, within a 1200 ms time window beginning at the onset of the second stimulus and extending to include the low frequency (e.g. alpha and beta) desynchronization / rebound effects. We averaged data across frequencies within each predetermined band (4-6 Hz for theta, 8-12 Hz for alpha, 14-30 Hz for beta) so as to obtain results specific to a band for ease of interpretation.

### EEG analysis: statistics

Our two main statistical analyses are outlined in Figure 2A. First, we investigated whether the AT “interaction” [(A+T)-(AT+N)] differed *across* the 7 asynchronies in a repeated-measures (i.e. dependent-samples) one-way rmANOVA (Figure 2A.1). Second, we also assessed whether the AT interaction was significant (i.e. different from zero) *within* each condition through a paired (i.e. dependent samples) t-test (Figure 2A.2) (i.e. contrasting (A+T) versus (AT+N) for each asynchrony). The rmANOVA and t-tests were performed separately for ERP, ITC, and TF power.

**Figure 2:**
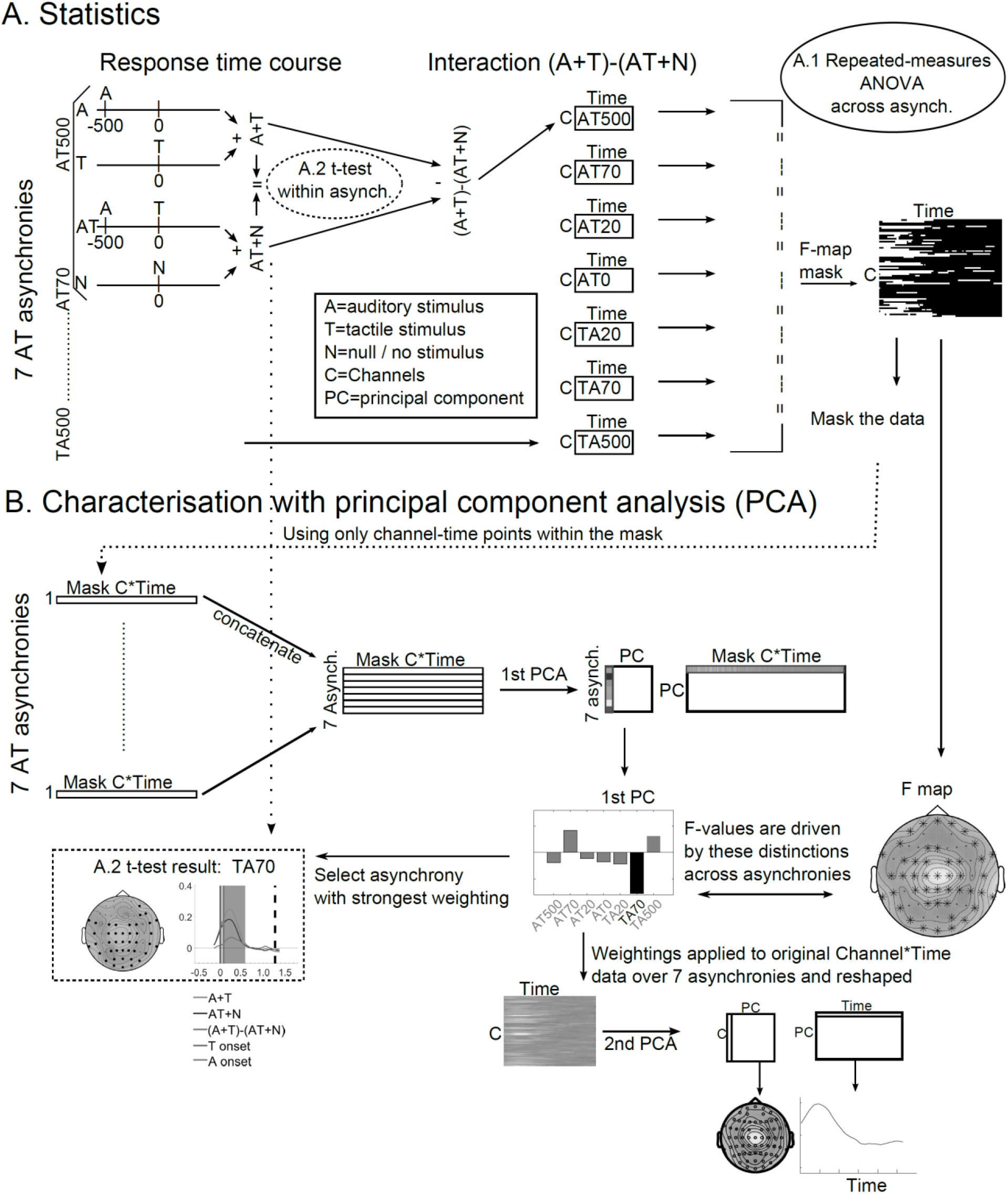
Overview of analyses stream. A. Statistics: within each asynchrony, the response time course (ERP, TF power, or ITC) for A alone (appropriately shifted in time to match a particular asynchrony) and T alone are summed, as are the time courses for AT (for a given asynchrony) and N. The difference of these, i.e. the audio-tactile “interaction” is computed for each asynchrony. A.1 These AT interactions computed for each asynchrony are compared across asynchronies in a repeated-measures (dependent-samples / paired) ANOVA (see Methods for details). A.2 Furthermore, the AT interaction within each asynchrony is tested with a paired (dependent samples) t-test. B. To assess the relative contributions of conditions to any significant modulatory effects of asynchrony on AT interaction in the rmANOVA as well as to relate the within-asynchrony assessment of AT interaction to the across-asynchrony assessment, two PCAs were sequentially performed. All AT interaction values from channel-time points in the significant cluster found in the rmANOVA were reshaped into a vector, one for each asynchrony, and then concatenated over asynchrony. This 7 X masked-channel-time matrix was entered in to the 1^st^ PCA, from which the first component was extracted and its weighting indicated the contribution of each asynchrony level to the effects revealed in the rmANOVA. This weighting across asynchrony levels was also applied to the original (non-masked) 7 X channel-time matrix to obtain the pattern across channel-time which reflects the differences across conditions. This vector was reshaped back to channels X time, and a 2^nd^ PCA was performed on this, from which the first component was extracted including the topography and time course. These are plotted alongside the within-asynchrony AT integration effect for the asynchrony which had the strongest condition weighting from the 1^st^ PCA.

To correct for multiple comparisons (over channels and time), we performed non-parametric cluster-based permutation tests for dependent (i.e. paired) samples, with the sum of the statistic (F or t-values) (i.e. max sum) across a cluster as cluster-level statistic and points for a cluster initially detected at an auxiliary uncorrected alpha threshold of 0.05 (Maris and Oostenveld 2007). All statistical results from power and ITC between 4-30 Hz were further corrected for testing over three frequency bands by dividing the p-value threshold by three (0.05 / 3 = 0.0167).

We illustrate a significant effect in a rmANOVA in a channel X time representation where all significant time points in a channel are highlighted (see Figure 2A, right and Figure 3A as an example). Further, we show an F-map where we show the topography of the F-values averaged across the time-window that includes time points where at least one channel was significant in the rmANOVA (see Figure 2B right and Figure 3B as an example).

**Figure 3.**
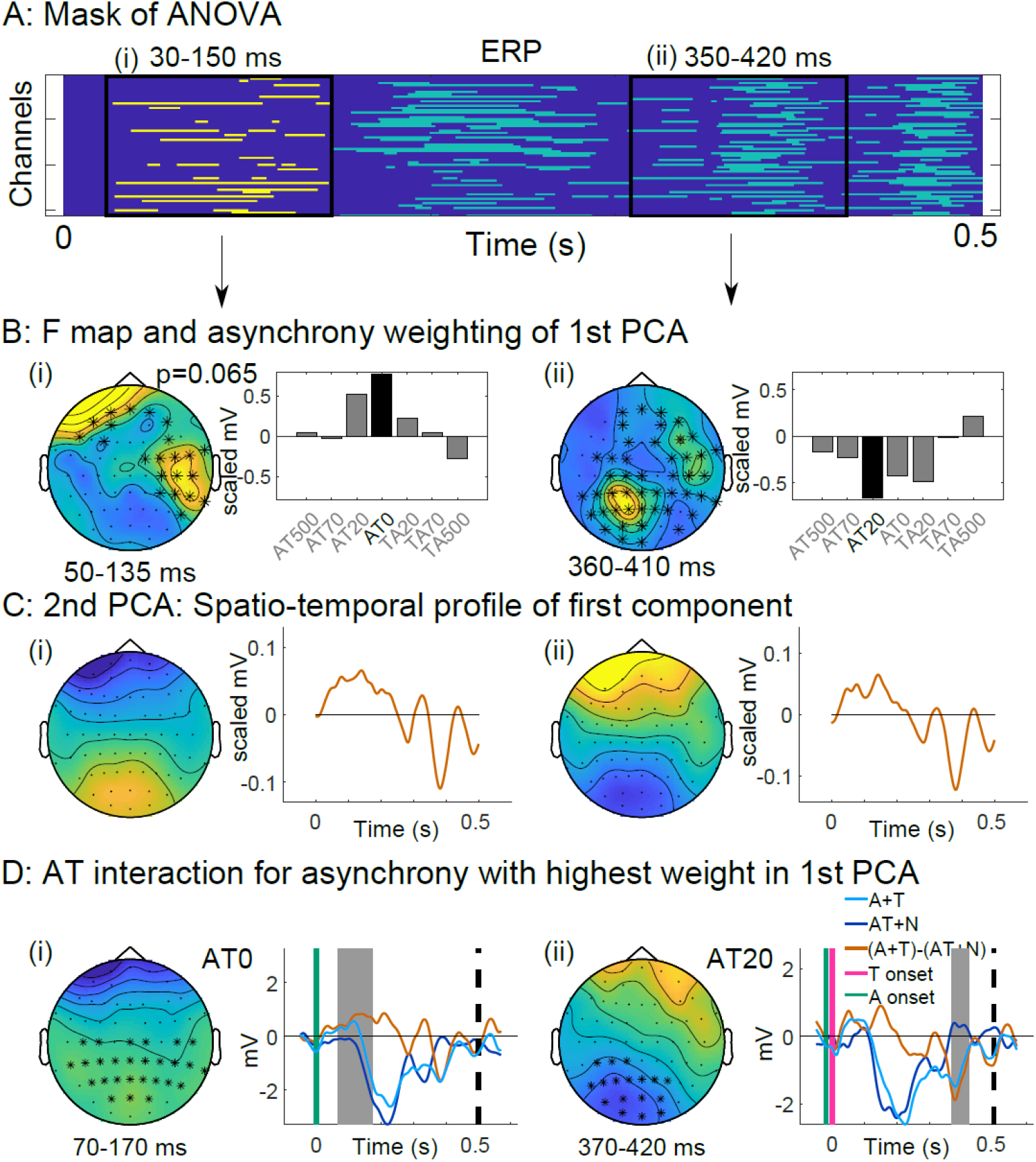
Early and late ERP effects. A. Channel X time matrix of points from the significant cluster in the rmANOVA test across asynchronies. For p-values, see Figure S2. Sub-clusters divided into time windows are shown separately in columns (i) and (ii) for the following. Time-window indicates cut-off for inclusion of masked points for the 1^st^ PCA. B. (left) topography of rmANOVA F values averaged from the time range indicated. This time window may be different than that used in A in order to avoid time points with only a few channels involved contributing to the average. (right) Relative weighting across asynchrony levels that contribute to this effect, according to PCA. C. A 2^nd^ PCA (spatiotemporal) was run to extract the strongest spatiotemporal profile contributing to the across-asynchrony differences; this topography and time course are shown. D. The AT interaction effect specific to one asynchrony (i.e. the one contributing most to the across-asynchrony weighting in sub-figure B) is shown, both for topography and time course (orange line depicts the AT interaction, as the difference of the A+T (light blue) and AT+N (dark blue); for more information on these, see Figure S3.

To further determine which asynchronies drive the significant main effect in the rmANOVA we characterised how it is expressed in the audiotactile interactions across asynchrony levels using a principal component analysis (PCA 1, see Figure 2B). For each asynchrony level we selected the contrast values of the AT interaction in each channel at the time points that were significant in the rmANOVA. The individual channel-time points *within* this rmANOVA cluster mask for each asynchrony were reshaped into a vector (one per asynchrony). These vectors were concatenated into a single matrix over asynchronies (7 asynchronies x Masked-channel-time-points). This matrix was entered into the PCA 1 that decomposes the matrix into a weighting (mixing) matrix that quantifies the expression over asynchrony of the principal components (PCs), which are in this case the representative masked-channel-time-point vectors; we focussed only on the first (strongest) PC as it is the one explaining the most variance. The first column of the weighting (mixing) matrix is a 7×1 vector indicating the contributing strength of each asynchrony AT to the main effect in the rmANOVA. For instance, in Figure 2B the 1^st^ PC indicates that the F-contrast from the rmANOVA is mainly driven by the TA70 asynchrony. In Figures 3-5, we then show the within-asynchrony interaction selectively for the asynchrony that received the greatest weight in the PC.

**Figure 4.**
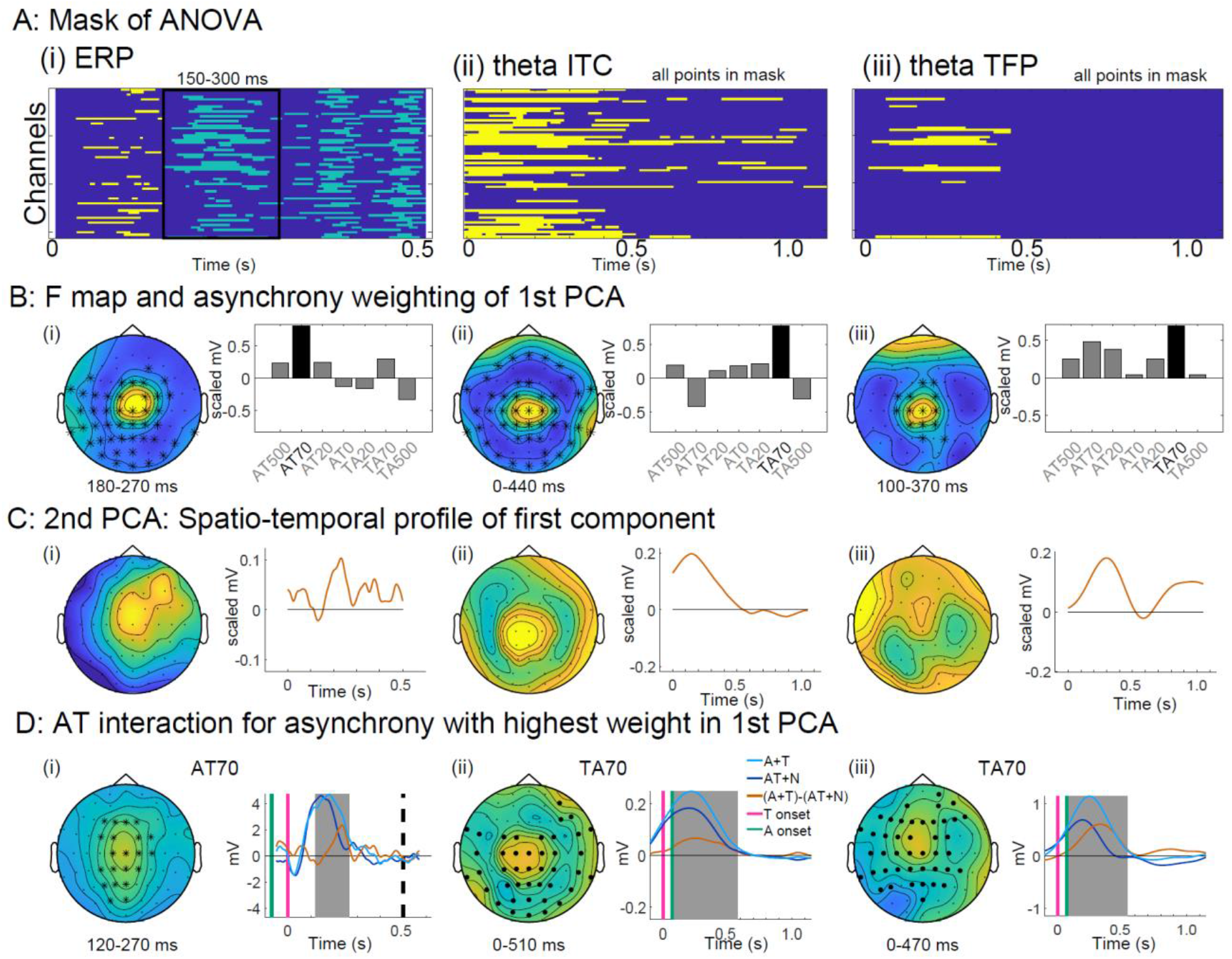
Same as for Figure 3, except that the columns are from the following data: (i) P200 ERP effect, (ii) theta-band ITC effect, and (iii) theta-band power effect. See also Figures S4, S6, and S7.

**Figure 5.**
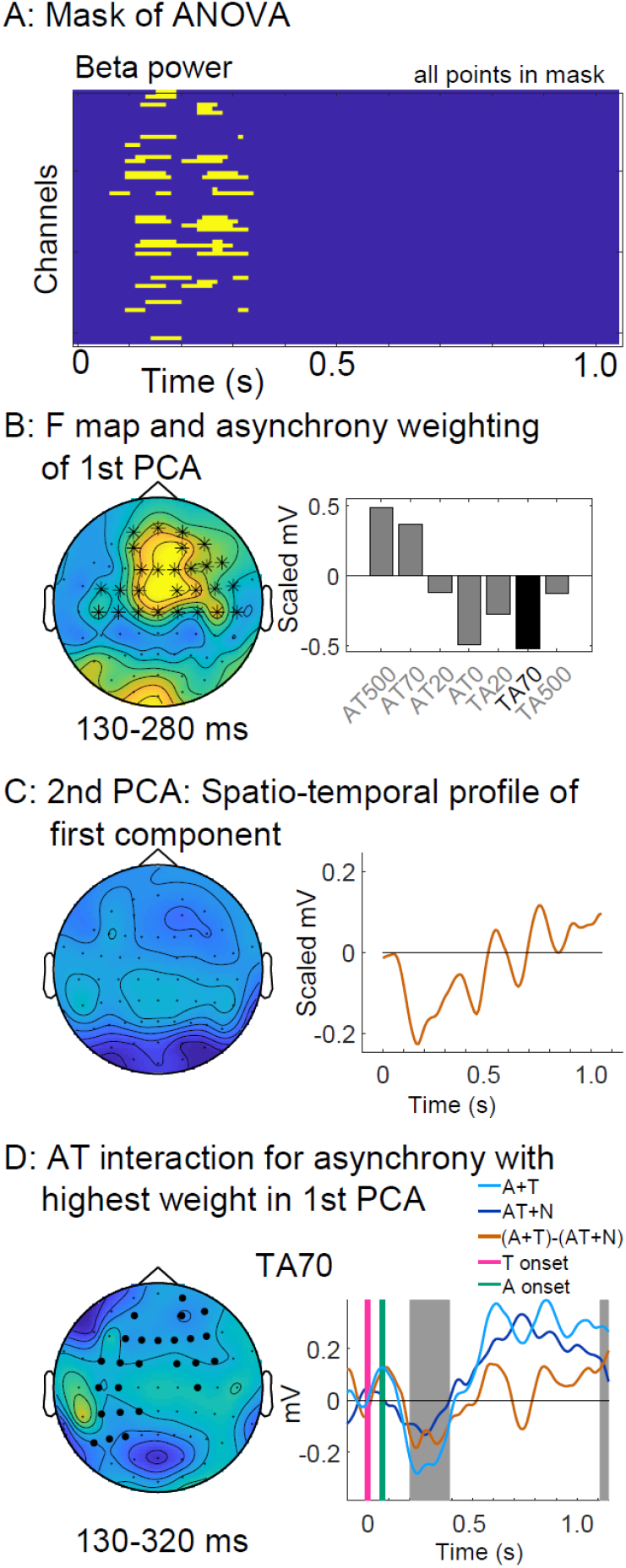
Same as for Figure 3, except the data are from the beta-band power effect. See also Figure S8.

Next, we characterised the spatiotemporal evolution of this effect by applying a 2^nd^ PCA. For this PCA 2, we linearly combined *all* channel-time points (not just those in the rmANOVA mask) of the audiotactile interactions across the 7 asynchrony levels (expressed as a matrix of size 7 X channel*time) using the weighting corresponding to the 1^st^ PC (7×1 vector) into a vector [of size 1 X channel*time]. We reshaped this vector back to the standard matrix [of size channel X time] and entered it in to the 2^nd^ PCA. We again focussed only on the first (i.e. strongest) PC and plotted its topography and time course (see Figure 2B, right bottom). We can then visually compare the topography and time course obtained from the 2^nd^ PCA with the time course of the audiotactile interaction effect for the asynchrony level that received the greatest weight in the PCA 1 (Figure 2B left). Note that any significant finding from the across-asynchrony rmANOVA, which indicates a difference in the AT interaction across asynchronies, need not necessarily appear as a significant finding in the within-asynchrony t-tests, which indicate a strong AT interaction effect for that one asynchrony, and vice versa. However, any correspondence found corroborates both the presence of the within asynchrony AT interaction and its dependence on asynchrony (i.e. significant difference in the AT interaction across asynchrony levels).

## Results

For the psychophysics study we report the redundant target effect as a behavioural index of audiotactile integration for each asynchrony level and contrasted across asynchronies. For the EEG data we assess how the multisensory interactions (AT+N ≠ A+T) for ERPs, inter-trial coherence (ITC), and time-frequency (TF) power differ across asynchrony levels in a rmANOVA. As described in detail in the methods section we then characterise the spatiotemporal profile of this interaction effect by applying a first and second stage principal component analysis. Moreover, we report the audiotactile interaction for the asynchrony level that mainly drives the effect in the rmANOVA as indicated by its weight in the first principal component analysis. For completeness and full characterisation of the data, the supplementary materials report all audiotactile interactions for each asynchrony level that was significant when tested independently in t-tests at a particular asynchrony level (i.e. at one of the seven levels of AT asynchrony: 0, ±20, ±70, and ±500 ms (Figure 1a).

### Behavioural results: reaction time facilitation tapered by TIW

As expected, we observed significantly faster (Figure S2 for p-values and t-values) response times for the AT relative to the fastest unisensory condition (i.e. *redundant target effect*) for asynchronies within a ≤ 70 ms window of integration (Figure 1b). Specifically, the RTEs (across subjects mean ± SEM) for the different asynchrony levels were: AT70 = 35 ms ± 6 ms, AT20= 38ms ± 5 ms, AT0 = 35ms ± 4 ms, TA20 = 33ms ± 4 ms, and TA70 = 24ms ± 4 ms. Surprisingly, we observed significantly slower response times for the AT500 relative to the unisensory auditory condition, i.e. a negative redundant target effect (across subjects’ mean ± SEM) = −16ms ± 4 ms.

Furthermore, we found that the redundant target effect differed significantly across the seven asynchrony conditions (one-way repeated-measures ANOVA (rmANOVA); F=25.4, df=6, p<1e-16). As a planned post-hoc test, we further found that, when restricting the conditions to those ≤ 70 ms asynchrony, the redundant target effect differed significantly across the five conditions within ≤ 70 ms (one-way rmANOVA; F=3.5, df=4, p=0.01). When restricting the comparison across conditions to those ≤ 20 ms, we did not find any significant difference (one-way rmANOVA; F=1.89, df=2, p>0.1).

In summary, our psychophysics study revealed that audiotactile interactions within a 70 ms temporal integration window (TIW) facilitate stimulus processing and response selection leading to faster response times. Furthermore, the response facilitation varied significantly across synchronies within 70 ms, while for near-synchronous stimuli within 20 ms they seem to be comparable.

### Audiotactile interactions for ERPs: limited to the behavioural TIW

Figure S3 shows the ERPs for the A, T, AT and N conditions. Both tactile-alone (pink) and auditory-alone (green) stimulation evoked a characteristic N100 followed by a P200, while the null condition is a flat baseline. The tactile and auditory stimulation together generate the AT evoked potentials across the different asynchrony levels (Figure S3, black). While the influences of both the tactile and auditory evoked responses are clearly visible in the AT responses, we can also observe small deviations from the unisensory responses. In the following, we investigate whether the “audiotactile interaction” ([AT+N]-[A+T]) significantly varies across asynchronies. If we observe a significant modulation of the AT interactions by asynchrony level in the rmANOVA, we then assess which asynchronies drive the effect based on the weights of the first PC. Further we report the AT interaction for the asynchrony level that is associated with the strongest weight in the PCA.

As shown in Figure 3, the rmANOVA across asynchronies for ERP revealed a near-threshold significant (p=0.065) cluster (shown in yellow) in an early time window (50-150 ms) (Figure 3A in yellow). Details of all statistical findings are in the table of Figure S2. The weights of the 1^st^ PC component (explaining 39.3% of the variance) (Figure 3B-i) indicate that this effect is mainly driven by AT interactions expressed for synchronous and to some extent also for near synchronous AT stimuli. Indeed, when testing for AT interactions separately for each asynchrony level we observed, for this early time window, a significant AT interaction only for AT synchronous stimuli (AT0) and trends also for the near synchronous stimuli ≤20 ms (i.e. AT±20 ms, see Figure S4). Further, Figure 3C and 3D indicate that the AT interactions in AT0 evolve during and after the N100 (70-170 ms), in both central and posterior sensors, with A+T being greater than AT+N.

The rmANOVA also revealed a significant temporally-extensive cluster (p=0.0005), spanning from about 150 ms to the end of the window tested at 500 ms (highlighted in light blue in the channel-time image in Figure 3A). This cluster included 3 ‘subclusters’ that were segregated in time, although linked together via a ‘bridge’ across channel-time space. Because the 3^rd^ subcluster can be attributed to eye artifacts based on it spatiotemporal and asynchrony profile, we do not discuss it further in the main manuscript (for completeness we show it in Figure S5).

The first subcluster extended from 180 to 270ms (Figure 4A). As shown in Figure 4A, it emerged with a similar spatiotemporal profile and was expressed across asynchrony levels similar to the audiotactile interactions observed for theta-band ITC and TF power; hence, it was plotted alongside these results. The F-values for the modulatory effects of asynchrony (i.e. rmANOVA) were most pronounced over frontocentral electrodes (Figure 4B-i). The first PC (explaining 31.9% of the variance) indicated that this effect was mainly driven by an audiotactile interaction at 70 ms asynchrony (AT70, Figure 4B (i) and Figure S4). Indeed, the spatiotemporal profile of the first PC of the 2^nd^ PCA (Figure 4C-i) and the audiotactile interaction for AT70 (Figure 4D-i) are very similar: they both emerge with a frontocentral topopgraphy and temporal peak around 200ms (see also Figure S4 for similar effect for TA70). As shown in Figure 4D-i, this AT interaction modulated the shape and magnitude of the P200: the P200 occurred earlier and was reduced in amplitude for the AT+N (dark blue) relative to A+T (light blue).

The second ‘sub-cluster’ (Figure 3A, ii) extended from 350 ms to 420 ms. The F-values for the modulatory effects of asynchrony (i.e. rmANOVA) were greatest over occipital electrodes. The weighting across asynchrony levels of the 1^st^ PCA (explaining 43.6% of the variance) indicates that this later audiotactile interaction effect was most pronounced for near-synchronous conditions ≤ 20ms and particularly for AT20. The further spatiotemporal characterisation of this effect (via 2^nd^ PCA) indicates a topography that varies from front to back and a time course with peak ∼400ms (Figure 3C-ii). This spatiotemporal profile was also found for the audiotactile interaction AT20 (Figure 3D-ii) and TA20 (Figure S4). Moreover, we note the similarities between the early (100ms) and late (400ms) effects in the ERP, both from rmANOVA (and subsequent PCA) and the within-asynchrony AT interactions.

In summary, we observed three distinct AT interaction effects for ERPs, all limited to AT asynchrony levels within the behavioural ≤ 70 ms TIW. The AT interactions at 100 ms and 400 ms were expressed mainly for synchronous and near synchronous AT stimuli. The AT interactions at 200 ms were mostly selective for 70 ms asynchrony and, as we will see in the next sections, related to AT interactions expressed in ITC and theta oscillatory power.

### Audiotactile interactions for ITC: selective for ±70 ms asynchronies

The across-asynchrony rmANOVA revealed that the AT interaction for the theta-band ITC differed significantly across the 7 asynchronies (p=0.0005; Figure 4A-ii and Figure S2 for statistics details). The topography of the maximal effect was also frontocentral (Figure 4B-ii) in line with the P200 ERP (Figure 4B-i). The 1^st^ PCA (explaining 35.9% of the variance) highlighted that the TA70 condition was the strongest driver (Figure 4B-ii). The 2^nd^ PCA revealed a spatiotemporal profile with a central topography most prominent around 200ms (Figure 4C-ii). Again this spatiotemporal profile was similar to the within-asynchrony audiotactile interactions (in this case, for +70 ms asynchrony, i.e. TA70), which also peaked at about 200 ms with a central topography (Figure 4D-ii) - thereby mimicking the AT interactions we observed for the P200 in the ERP analysis (Figure 4i). As shown in the supplementary results (Figure S6) we also observed a similar AT interaction effect for −70 ms asynchrony (i.e. AT70). Surprisingly, as seen in both the PCA weightings across asynchrony levels (Figure 4B-ii) as well as within-asynchrony multisensory integration effects (Figure S6), the summed ‘A+T’ ITC was *smaller* than the summed ‘AT+N’ for the AT70, but *greater* for tactile leading TA70 condition. Thus, the direction of the audiotactile theta-band ITC interaction depends on whether the auditory or the tactile sense is leading. No significant ITC results were found for alpha, beta, or gamma bands. In summary, the AT interactions for the theta-band ITC were most prominent for 70 ms asynchronies and most likely associated with the ERP effects at the same post-stimulus latency and asynchrony conditions.

### Audiotactile interactions for time-frequency power across AT asynchronies

#### Theta power

The across-asynchrony rmANOVA revealed a marginally significant (see Methods) single cluster (p=0.024; Figure 4A-iii) primarily with frontocentral topography (Figure 4B-iii) and strongest from 200-300 ms (Figure 4A-iii). The 1^st^ PCA (explaining 53.5% of the variance) showed that TA70 (and AT70) mainly drove this difference in audiotactile interactions across asynchrony levels (Figure 4B-iii). The 2^nd^ PCA showed a spatiotemporal profile of this 1^st^ principal component (Figure 4C-iii) with a frontocentral topography peaking around 200-300 ms. Likewise, the audiotactile interaction for the TA70 asynchrony level emerged with a frontocentral topography peaking at about 200-300 ms (Figure 4D-iii; see Figure S2 for statistics and also Figure S7 for other asynchronies). These frontocentral AT interactions arose as a result of the AT+N power peak being weaker and decaying earlier relative to the A+T sum. Figure 4 also highlights the point that the audiotactile interactions for the P200 ERP, the theta ITC and the theta TFP emerge with a similar spatiotemporal profile. Further they are all most pronounced for ±70 ms asynchrony.

#### Beta power

The rmANOVA revealed significant differences in audiotactile interactions in beta power across asynchronies (p=0.001; Figure S2) in an early (100-350 ms) cluster (Figure 5A) with frontocentral topography (Figure 5B). The weighting from the 1^st^ PCA (explaining 38.5% of the variance) shows that AT70 and TA70 asynchronies contributed to this difference in opposite directions (Figure 5B). These results again suggest that not only the absolute asynchrony but also the direction is a critical determinant of audiotactile interactions. In contrast to the asynchrony-specific weights observed in the other ERP, ITC and TFP analyses, we did not observe one single asynchrony that selectively contributed to the significant difference across asynchrony levels. Instead, AT70, TA70, AT0 and AT500 jointly drove the effect observed in the rmANOVA (Figure 5B). Not surprisingly the spatiotemporal profile associated with the 1^st^ principal component (Figure 5C) and the TA70 asynchrony (Figure 5D) that formally received the greatest weight are therefore slightly less in correspondence. Although the time courses of the first PC (Figure 5C) and the AT interaction for the TA70 asynchrony (Figure 5D) roughly match (keeping in mind that time=0 in Figure 5C means the onset of the second stimulus) and both emerged with a predominantly central topography, the correspondence is not as striking as for the previous results. As shown in Figure 5D, the audiotactile interaction in beta oscillatory power for TA70 relied on a stronger suppression in beta power (event-related desynchronization; ERD) for A+T than AT+N stimuli around 250 ms post-stimulation followed by a rebound (event-related synchronisation; ERS) above and beyond baseline, around 800-1000 ms post-stimulation (for AT interactions at other asynchronies see Figure S8). In summary, audiotactile interactions expressed in beta power differed at about 200 ms across AT asynchronies, depending not only on the absolute level of AT asynchrony, but also whether the auditory or tactile signal were leading.

#### Alpha and gamma power

The rmANOVA across asynchronies did not show any significant differences in the alpha or gamma band.

## Discussion

The current study presented A, T, and AT stimuli at several asynchrony levels to investigate the temporal constraints that govern behavioural response facilitation and neural AT interactions for ERPs, ITC, and induced TF power.

Consistent with previous research (Colonius and Diederich 2004), we observed an inverted U-shape function for the behavioural AT benefit – also coined the redundant target effect (Miller 1982) – that was maximal for synchronous AT combinations and tapered off with increasing AT asynchrony within a TIW of ≤70 ms (Figure 1B) (Zampini et al. 2005).

At the neural level we observed early AT interactions for evoked responses (ERP) at about 110 ms post-stimulus (Figures 3 and S4), which dovetails nicely with previous research showing multisensory modulations of the N1 auditory component by visual and tactile stimuli (Foxe et al. 2000; Lutkenhoner et al. 2002; Murray et al. 2005; Sperdin et al. 2009; Stekelenburg and Vroomen 2009). Critically, our observed early AT interactions were sensitive to the relative timing of the AT stimuli: they were most pronounced for synchronous AT stimuli and tapered off within a small TIW of ≤20 ms (Figure 3B-i. weights across asynchrony). This temporal precision may be enhanced for interactions of tactile with other sensory signals, because tactile latencies are fixed for a particular body location and do not vary depending on the distance of the stimulus from the observer as in audition and vision. The short latency and narrow temporal binding window points towards neural interactions in low level or even primary auditory cortices that may rely on direct connectivity between sensory areas (Fu et al. 2003; Cappe and Barone 2005; de la Mothe et al. 2006a; Smiley et al. 2007) or thalamic mechanisms (de la Mothe et al. 2006b; Hackett et al. 2007; Cappe et al. 2009) and that increase the saliency of AT events leading to faster and more accurate detection.

Later, at about 400 ms post-stimulus, we observed audiotactile ERP interactions that were again most pronounced for synchronous AT stimuli and confined to the TIW of ≤20 ms (Figures 3-ii and S4). These later interactions may reflect top-down modulatory neural processes in lower regions via feed-back loops (Falchier et al. 2002; Schroeder and Foxe 2002; Clavagnier et al. 2004). The expression of both early and late ERP interactions followed a U-shape function (Figure 3B-i and 3B-ii) thereby mimicking the asynchrony profile of the redundant target effect that characterised observers’ behaviour.

While the ERP effects at ∼125 ms and ∼400 ms post-stimulus were constrained by classical temporal integration windows, the AT interactions for the P200 ERP component were most pronounced for ± 70 ms AT asynchrony and absent for near-synchronous AT stimulation (see Figures 4B-i and S4). Both the auditory and the tactile unisensory P200 are thought to be generated in regions previously implicated in audiotactile integration (Foxe et al. 2002; Kayser et al. 2005; Murray et al. 2005; Schurmann et al. 2006) such as the auditory belt area CM or planum temporale (Godey et al. 2001; Crowley and Colrain 2004; Smiley et al. 2007) and secondary somatosensory areas (Forss et al. 1994; Disbrow et al. 2001), respectively. Our results show that AT integration facilitates neural processing at about 200 ms post-stimulus: the P200 peaks earlier, is smaller, and/or decays faster for the AT+N sum when compared to the sum of the unisensory A and T conditions (Figures 4D-i and S4), consistent with multisensory literature, e.g. (Rowland et al. 2007).

The P200 ERP effects were directly related to AT interactions for theta-band ITC and TFP that emerged with a central topography again at ∼200 ms post-stimulus primarily for ± 70 ms AT asynchrony (Figure 4). Critically, whilst the ERP and theta-band power interactions followed a similar temporal profile and topography irrespective of whether the auditory or the tactile stimulus is leading, the ITC effects were inverted for auditory relative to tactile leading stimulation (i.e., the condition weighting for P200 and theta power are in the same direction for AT70 and TA70, whereas the condition weightings for these asynchronies are opposing for theta ITC; Figure 4B). This dissociation between ERP and ITC is mathematically possible and one possible mechanism can be shown to produce this effect in simulation (https://github.com/johanna-zumer/audtac/blob/master/simulate_70results.m). The selectivity of the P200 and the phase coherence effects for ± 70 ms AT asynchrony may be best accounted for by mechanisms of phase resetting that have previously been implicated in audiotactile and audiovisual interactions in auditory cortices (Lakatos et al. 2007; Kayser et al. 2008; Thorne et al. 2011). From a functional perspective, a preceding tactile stimulus may reset the phase in auditory cortices and thereby facilitate the localization of an auditory stimulus that is presented 70 ms later. Likewise, a preceding auditory stimulus may provide an alert to facilitate tactile processing and possible avoidance actions. Not only have tones been shown to elicit responses in somatosensory cortex (Borgest and Ermolaeva 1975; Liang et al. 2013), but also an *inhibitory* multisensory interaction by auditory stimulation was found in cat somatosensory area SIV (Dehner et al. 2004) and auditory projections were found to inhibitory interneurons in cat SIV (Keniston et al. 2010). In summary, our P200 ERP and theta-band ITC and power results are supported by evidence of bidirectional audiotactile integration, especially to association cortices, and of directional asymmetries in the AT interaction (Cecere et al. 2017).

The AT interactions discussed so far were moulded by two distinct neural mechanisms: (i) ERP effects at ∼100 and ∼400 ms that followed a U-shape function across asynchronies, mimicking the temporal binding window at the behavioural level and (ii) effects primarily driven by the ± 70 ms AT asynchronies, reflected in the P200, theta ITC, and theta power, and may be mediated by mechanisms of phase resetting.

In contrast, AT interactions for induced beta oscillatory power were expressed across a set of asynchrony levels. As shown in Figure S8 (bottom row), both auditory and tactile stimuli suppressed beta oscillatory power (event-related desynchronization; ERD) at about 200-400 ms, related to a release from inhibition, followed by a rebound in power beyond baseline levels from about 600 ms – 1200 ms post-stimulus (event-related synchronisation; ERS), related to resetting and recovery (Pfurtscheller and Lopes da Silva 1999; Neuper and Pfurtscheller 2001). Beta-band AT interaction effects differed significantly across asynchronies only in the early (100-300 ms) time window (Figure 5). In the late (∼1000 ms) window we observed AT interactions in the beta rebound for a few specific asynchronies, but no significant difference across asynchronies (Figure S8 and discussed further in the supplementary discussion). The early effects were supported by several asynchronies and indicated a possible dependence on which sense came first, as the weighting across conditions (Figure 5B) flipped for A-leading versus T-leading. This is consistent with other studies that have shown sense-leading dependencies (e.g. Cecere et al., 2017) although novel here for beta-band effects. In contrast to the AT interactions in ERP, ITC and theta TF power the AT interactions for beta power were not limited to the ± 70 ms temporal integration window but included the AT500 condition. The AT interactions for beta power may thus generally reflect non-specific mechanisms of multisensory priming or attention by which a preceding A (or T) signal may alert the observer to imminent touch (or sound) events, in light of the debate as to whether cross-modal stimuli with asynchronies up to 500-600 ms may be actually *integrated* or whether the first stimulus (only) primes and/or draws exogenous (spatial) cross-modal attention (Macaluso et al. 2001; McDonald et al. 2001; Stein et al. 2010). Alternatively, the AT interactions for beta power may rely on several mechanisms depending on the AT asynchrony level, in which case the topography shown in Figure 5B and 5C may reflect the combination of these three (or more) mechanisms.

Whilst this study did not directly compare the dual AT conditions against dual AA or TT conditions, it has been shown (Forster et al. 2002) that the reaction times to visual-tactile dual stimuli were faster than for dual tactile or dual visual, indicating that the redundant target effect reflects special multisensory processing above and beyond that of integrating two stimuli (of the same sense). In the same way, regarding the EEG data, we did not explicitly contrast multisensory against dual unisensory conditions (e.g. AT+AT versus AA+TT) and thus are not accounting for general neural mechanisms of processing two stimuli versus one at a time. However, we argue that by contrasting different multisensory asynchrony conditions against each other (i.e. the rmANOVA applied to both reaction time and EEG data), we mitigate this interpretational ambiguity.

To conclude, this psychophysics-EEG study unravels a multitude of neural interactions, which arose with different temporal constraints: interactions were confined to a narrow TIW for early and late ERPs, interactions were specific for one particular asynchrony for a middle ERP, theta ITC and theta power, and beta-band power modulations indicated both early and late (rebound) effects. This diversity of temporal profiles demonstrates that distinct neural mechanisms govern a cascade of multisensory integration processes.

## Acknowledgements

We thank Christoph Braun, Jürgen Dax, and Elisa Leonardelli for assistance with the tactile device and Máté Aller for assistance with EEG and stimulus setup.

