## Supplementary Information for "Distinct neural mechanisms and temporal constraints govern a cascade of audiotactile interactions"

**Supplementary Results and Discussion**

The results presented in the main manuscript asked the question whether audiotactile interactions differed across asynchrony levels. Hence, it focused on the rmANOVA results across asynchronies and characterised the results using subsequent PCA analysis. Further, we presented the audiotactile interaction for the asynchrony level that contributed most to the significant difference across asynchronies.  For complete characterisation of our data, the following supplementary results present all individual asynchrony results (i.e. paired t-test results).  Please note that these individual analyses are complementary to our rmANOVA. For instance, an AT interaction that is significant only for one particular AT asynchrony level does not imply that the AT interactions are significantly different across asynchronies and vice versa.

An example of the light reflectance pattern from the tactile device is shown, indicating the evidence we have to confirm that the tactile device touched the skin for ~200 ms (Figure S1). Please note the location of absolute ‘0 ms’ on this figure’s scale is arbitrary; each trial is re-defined so that time=0 is at the tactile onset.

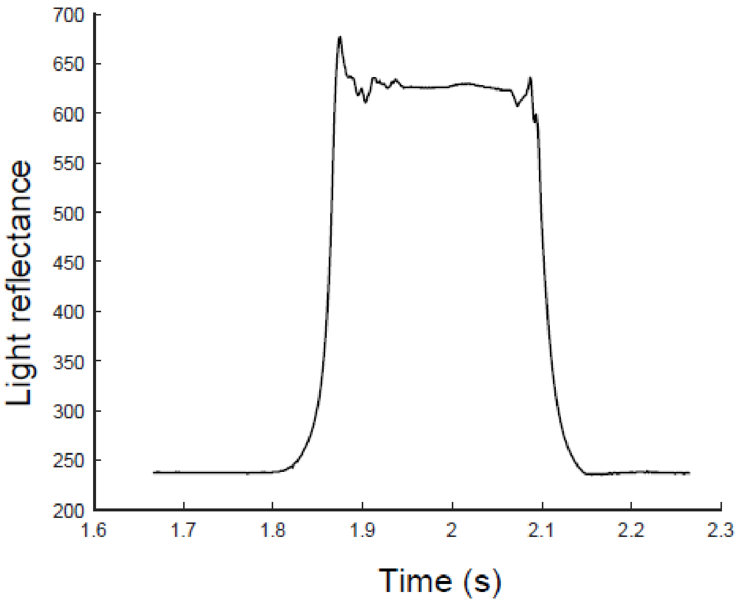

**Figure S1:** Output from one example trial of the light reflectance measured from the tip of the probe used for tactile stimulation. There is a small over-shoot at both the beginning and end of the 200 ms duration stimulation.

To provide an overview of all significance results, Figure S2 lists the p-values from the rmANOVA and paired t-tests systematically in a table organized in different time windows.

| MSI effects | Behaviour: | Neural Post-stimulus Latency: | | | |
|---|---|---|---|---|---|
| Asynchrony: | RTE | ~125 ms | ~200 ms | ~400 ms | ~800-1100 ms |
| **ANOVA across asynchronies** | p<1e-16<br>F=25.4<br>(df=6) | *ERP 30-150 ms (p=0.065)* | ERP 150-300 ms (p=0.0005)<br>θ ITC 0-500+ ms (p=0.0005)<br>*ϑ TFP 40-440 ms (p=0.0235)*<br>β TFP 70-340 ms (p=0.001) | | |
| AT500 | p=0.001<br>t=-3.7 | | *ϐ ITC 190-270 ms (p=0.0465)* | θ TFP 150-1200 ms (p=0.0005) | *ϐ TFP 790-950 ms (p=0.035)* |
| AT70 | p=1.2e-5<br>t=5.7 | | ERP 120-270 ms (p=0.010 )<br>θ ITC 0-260 ms (p=0.0005)<br>θ TFP 100-470 ms (p=0.0005) | | β TFP 1040-1150 ms (p=0.0115) |
| AT20 | p=7.2e-8<br>t=8.1 | *ERP 40-100 ms (p=0.072)* | θ ITC 40-390 ms (p=0.011)<br>θ TFP 50-420 ms (p=0.005) | ERP 370-420 ms (p=0.022 | *ϑ TFP 890-1200 ms (p=0.046)*<br>*α TFP 750-960 ms (p=0.0455)*<br>β TFP 870-1060 ms (p=0.0155) |
| AT0 | p=1.1e-7<br>t=7.8 | ERP 70-170 ms (p=0.017) | *ϑ ITC 0-320 ms (p=0.036)* | *ERP 370-420 ms (p=0.28)* | β TFP 860-1120 ms (p=0.0015) |
| TA20 | p=2.1e-8<br>t=8.7 | | *ϑ ITC 0-270 ms (p=0.0395)*<br>*ϑ TFP 90-390 ms (p=0.048)*<br>*ϐ TFP 200-300 ms (p=0.0245)* | ERP 340-400 ms (p=0.03) | |
| TA70 | p=8.8e-5<br>t=4.8 | | ERP 140-220 ms (p=0.007)<br>θ ITC 0-510 ms (p=0.001)<br>θ TFP 0-470 ms (p=0.0005)<br>β TFP 130-320 ms (p=0.007) | | α TFP 980-1200 ms (p=0.0075)<br>*ϐ TFP 1040-1110 ms (p=0.045)* |
| TA500 | | | θ TFP 0-400 ms (p=0.0075) | | *ϑ ITC 800-950 ms (p=0.019)* |

**Figure S2**: Statistics for behavioural and neural results (columns) for each AT asynchrony and rmANOVA across asynchrony (rows). *Behavioural redundant target effect (RTE) (left-most data column)*: rmANOVA comparing the RTE across asynchronies and paired t-tests (sample size: N=22; degrees of freedom = 21) comparing the AT response time with the minimal unisensory response time. For AT500 the AT response was slower than the minimal unisensory response (negative t-value). The “e” indicates “x 10^”. *Neural AT interactions* [(A+T) – (AT + N)] for ERPs (blue), ITC (violet), and TFP (red) listed in separate columns for different latency ranges. Non-parametric permutation rmANOVA of the AT interactions comparing across asynchronies (first row) and non-parametric permutation dependent/paired samples t-tests (sample size N=22) comparing A+T with AT+N (subsequent rows). The p-values are reported at the cluster level (max sum) corrected for multiple comparisons over channels and time. The ERP significance is set to a threshold of $p<0.05$; the time-frequency results (ITC and TFP) are further corrected for the 3 frequency bands (theta, alpha, and beta) and so are subject to a threshold of $p<0.017=0.05/3$. P-values in italics indicate a non-significant trend.

To supplement Figure 1C of the main text, Figure S3 shows the ERPs for all seven asynchrony conditions and in colour.

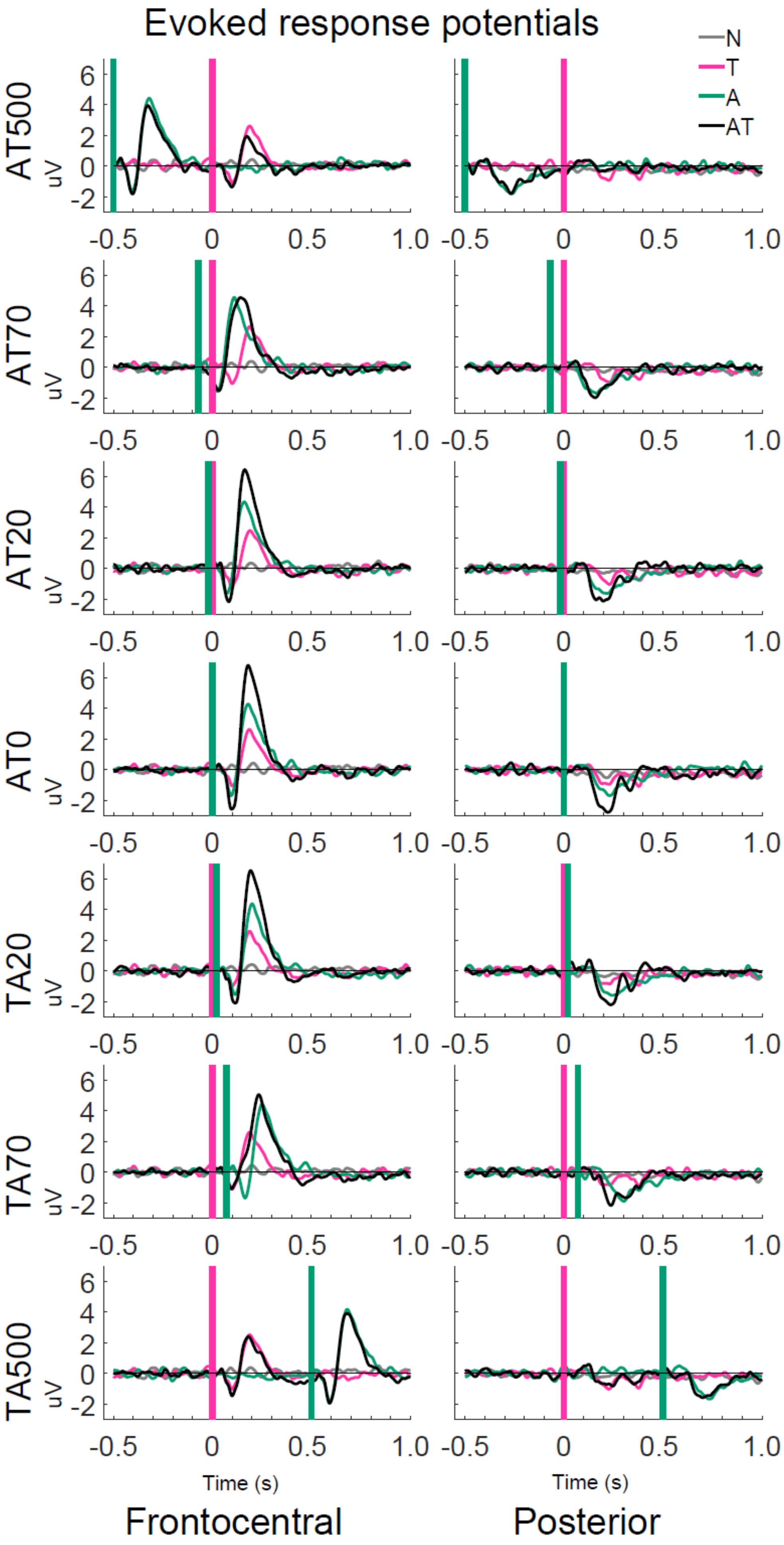

Evoked response potentials
N
T
A
AT
AT500
AT70
AT20
AT0
TA20
TA70
TA500
uV
6
4
2
0
-2
-0.5
0
0.5
1.0
Time (s)
Frontocentral
Posterior

**Figure S3.** Evoked response potentials for N, A, T, and AT conditions for frontocentral ['Fz' 'Cz' 'F1' 'F2' 'FC1' 'FC2' 'C1' 'C2'] and posterior ['CP5' 'POz' 'Pz' 'P3' 'P4' 'C4' 'O1' 'O2' 'P7' 'PO7'] sets of sensors. The 'A' evoked response is shifted by the appropriate asynchrony to align with the auditory onset in the corresponding AT condition.

**Audiotactile interactions for ERPs: limited to the behavioural TIW**

Figure S4 shows the ERPs for the sum over A+T (dark blue), sum over AT + N (light blue), and the difference (A+T) – (AT + N), i.e. the audiotactile interaction effects across different asynchrony levels. For ERPs we observed three AT interaction effects that differed in their expression across levels of AT asynchrony.

The first AT interaction effect arose early, at about 100 ms post-stimulus, with a central topography and was significant only for the synchronous condition (Figure S4, AT0 row). Specifically, a modulation, during and after the N100 (70-170 ms), was found in both central and posterior sensors, with the A+T greater than the AT+N during this time. We note that a trend for this spatiotemporal effect was also observed for the AT20 condition.

The second AT interaction effect emerged at about 200 ms after the second stimulus (latency range: 140-220 ms), was most pronounced over frontocentral electrodes, and was selective for the asynchrony of ±70 ms (Figure S4, AT70 and TA70 rows). This AT interaction modulated the shape and magnitude of the P200: the P200 occurred earlier and was reduced in amplitude for the AT+N relative to A+T.

The third AT interaction effect, where A+T was more negative than the AT+N, arose later at about 370-400 ms mainly over posterior electrodes for AT asynchrony conditions within a ≤ 20 ms temporal integration window (Figure S4, AT20, AT0, and TA20 rows). Even though

71 this AT interaction effect was significant only for AT20 and TA20, we observed a qualitatively similar pattern for the synchronous AT0 condition.

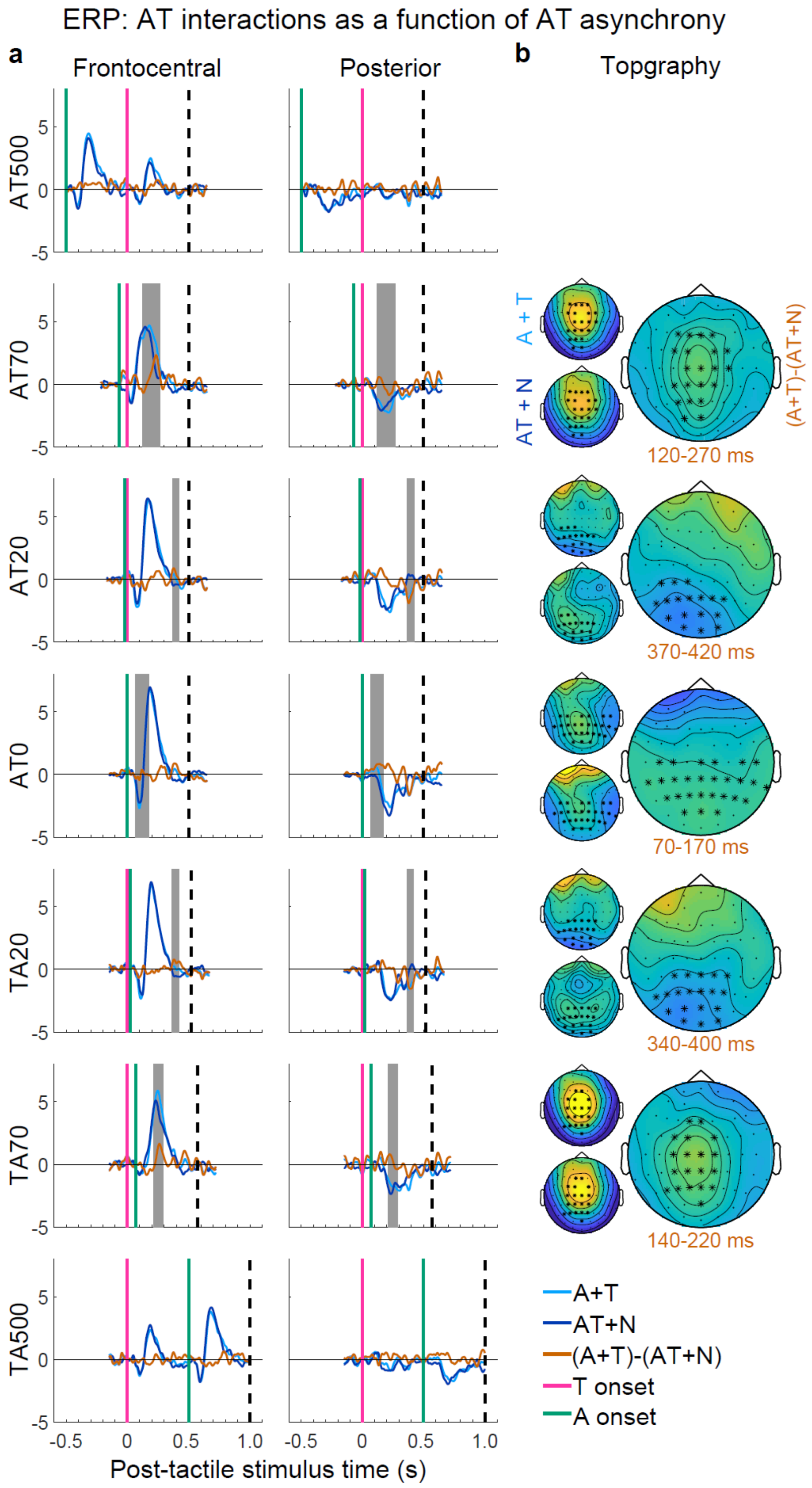


73

**Figure S4:** Evoked response potentials. Each row shows the audiotactile interaction for a particular level of AT asynchrony. (A) ERPs of the sum of the auditory and tactile (A+T; light blue), the sum of audiotactile plus null (AT+N; dark blue), and the audiotactile interaction, i.e. the difference ([A+T]-[AT+N], orange). Green = auditory onset, pink = tactile onset. Shaded grey areas indicate the timing of significant AT interactions at $p < 0.05$ corrected at the cluster level for multiple comparisons across electrodes and time points within a 500 ms window starting with the second stimulus and limited by the black dashed line. (B) Topographies of the sums: A+T, AT+N, and (A+T)-(AT+N) for time windows of significant AT interactions. The time windows written in orange are relative to the onset of the second stimulus. A black star over an electrode indicates that it is part of a significant cluster.

A fourth 'peak' in the rmANOVA (440-500 ms) did not correspond to any within-asynchrony effects (paired t-test results) in the ERP; however, we show the rmANOVA and PCA analyses in Figure S5 for completeness and to illustrate that it is likely related to an eye artefact specific to the TA500 condition, rather than a relevant neural finding.

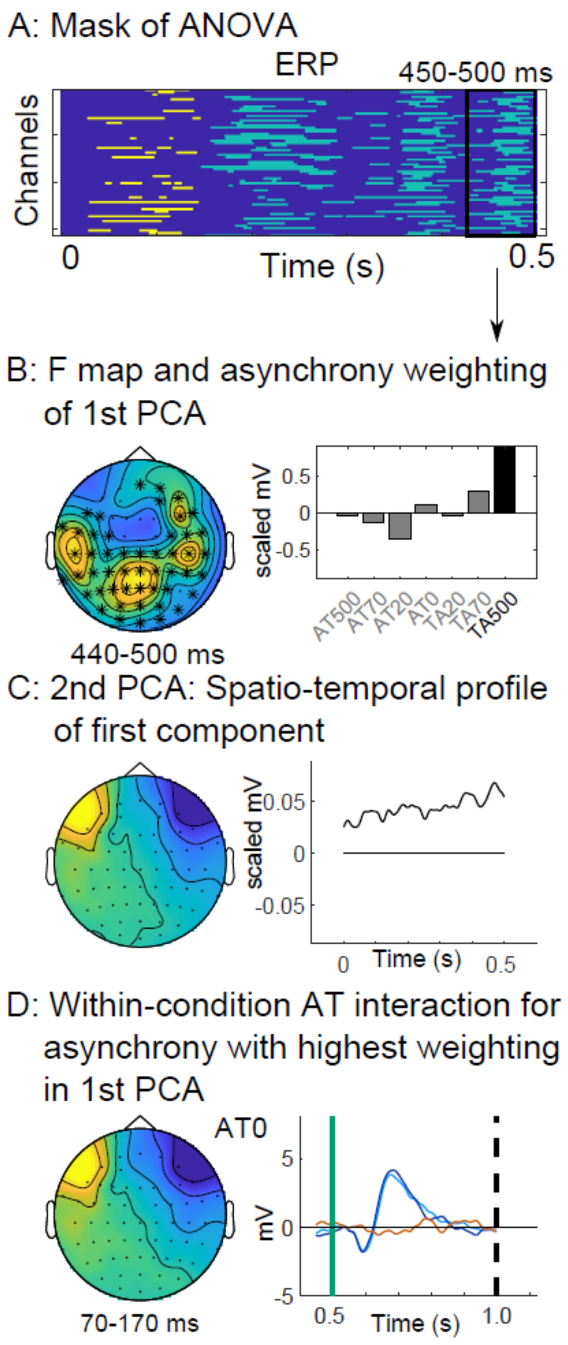


**Figure S5:** Similar to main Figure 3, for final/late sub-cluster in ERP from rmANOVA.

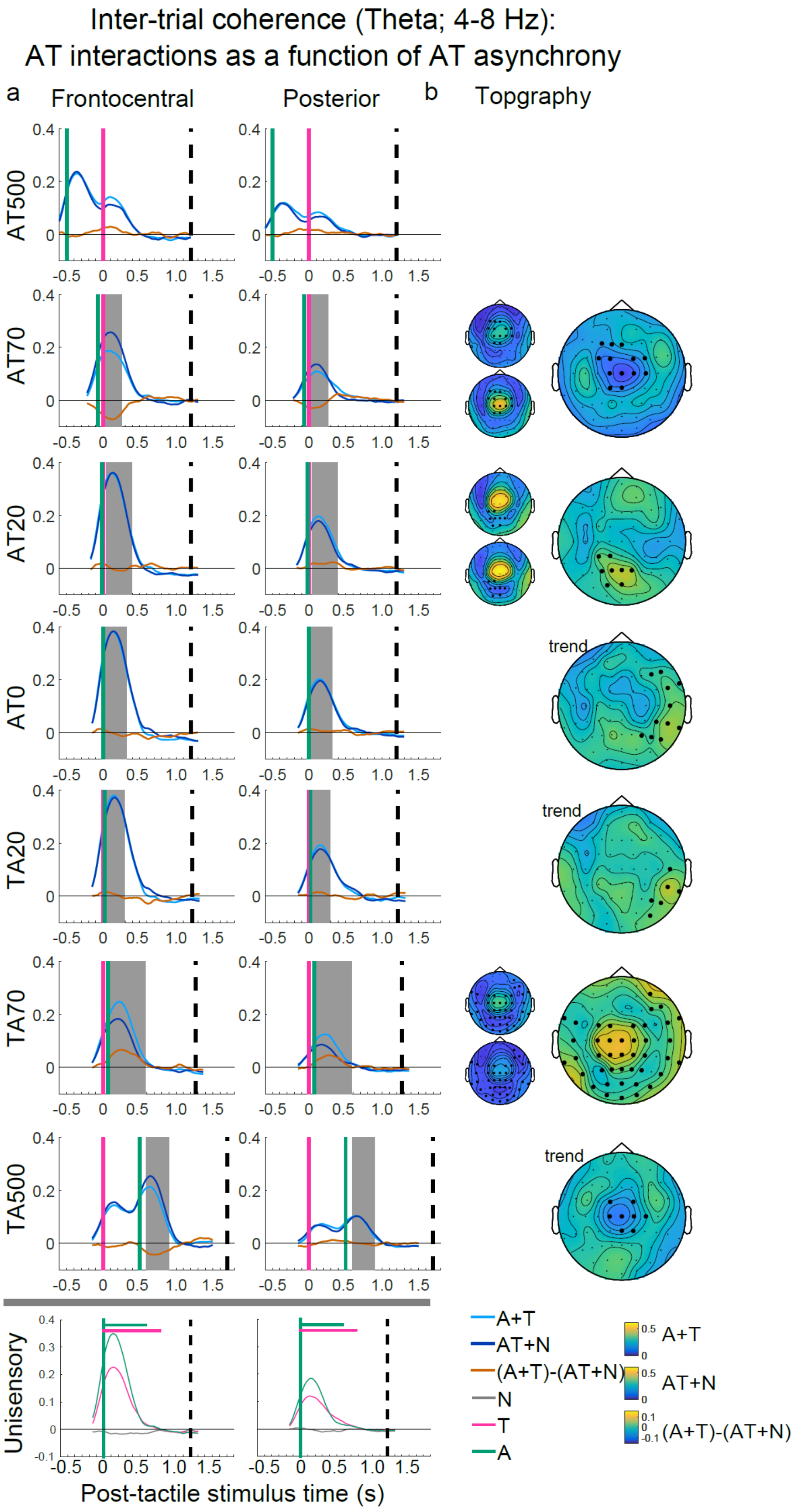
Inter-trial coherence (Theta; 4-8 Hz):
AT interactions as a function of AT asynchrony
a
Frontocentral
Posterior
b
Topgraphy
AT500
AT70
AT20
AT0
TA20
TA70
TA500
Unisensory
trend
A+T
AT+N
(A+T)-(AT+N)
N
T
A
Post-tactile stimulus time (s)

**Figure S6:** Inter-trial coherence. Each row shows the audiotactile interaction in the ITC for a particular level of AT asynchrony, plus the unisensory conditions. (A) ITC of the sum of the auditory and tactile (A+T; light blue), the sum of audiotactile plus null (AT+N; dark blue), and the audiotactile interaction, i.e. the difference ([A+T]-[AT+N], orange). The bottom row shows the Null (grey), Tactile (pink), and Auditory (green) conditions. Green = auditory onset, pink = tactile onset. Shaded grey areas indicate the timing of significant AT interactions at $p < 0.05$ corrected at the cluster level for multiple comparisons across electrodes and time points within a 1200 ms window starting with the second stimulus and limited by the black dashed line. Note that these effects were further corrected for 3 frequency bands (theta, alpha, beta) and are subject to a final threshold of $p<0.017$; any result with p-value between 0.017 to 0.05 is labelled as a 'trend'. (B) Topographies of the sums: A+T, AT+N, and (A+T)-(AT+N) for time windows of significant AT interactions. The time windows written in orange are relative to the onset of the second stimulus. A black star over an electrode indicates that it is part of a significant cluster.

**Audiotactile interactions for ITC: selective for ±70 ms asynchronies**

Figure S6 shows the ITC for the sum over A+T (light blue), sum over AT + N (dark blue), and the difference (A+T) – (AT + N) (orange), i.e. the audiotactile interaction effects across different asynchrony levels, as well as unisensory and null conditions separately. We observed significant AT interactions for ITC in the theta band (4-8 Hz) specifically for ±70 ms asynchrony levels (Figure S6, AT70 and TA70 rows; Figure S2 for significance test results). As shown in Figure S6, the summed 'AT+N' ITC was greater than the summed 'A+T' for the auditory leading AT70, but smaller for tactile leading TA70 condition. Thus, the direction of the audiotactile ITC interaction depends on whether the auditory or the tactile sense is leading. The AT interaction arose at about 200 ms post-stimulus and was most prominent over frontocentral electrodes, mimicking the AT interactions we observed for the P200 in the ERP analysis (Figure S3, AT70 and TA70 rows). In summary, the AT interactions for the theta-band ITC were selective for ±70 ms asynchronies and most likely associated with the ERP effects at the same post-stimulus latency and asynchrony conditions.

**Audiotactile interactions for time-frequency power across AT asynchronies**

Figures S7 and S8 show the TF power (for theta and beta bands, respectively) for the sum over A+T (light blue), sum over AT + N (dark blue), and the difference (A+T) – (AT + N) (orange), i.e. the audiotactile interaction effects across different asynchrony levels, as well as unisensory and null conditions separately. For significance test results, see Figure S2.

*Theta power*: Both auditory and tactile stimuli induced theta power peaking at about 200 ms post-stimulus primarily over fronto-central electrodes (Figure S7; bottom Unisensory row). This peak in theta power corresponds to the P200 (Figure S4) in the ERP analysis and an increase in ITC (Figure S6). Note that our data illustrate the point that the 'A+T' sum (Figure S7: AT0 light blue), which was computed by first summing trials before frequency transformation according to Senkowski et al. (2007), is indeed different than if the power of the tactile (Figure S7: Unisensory pink) and auditory (Figure S7: Unisensory green) had first been computed and then summed.

We observed significant AT interactions in the theta band at about 200 ms post-stimulus over fronto-central electrodes across several asynchrony levels including AT70, AT20, and TA70. These fronto-central AT interactions arose as a result of the AT+N power peak being weaker and decaying earlier relative to the A+T sum. Critically, these fronto-central AT interactions for theta power were most pronounced for ± 70 ms asynchrony levels, expressed less strongly for ± 20 ms and ± 500 ms asynchrony and completely absent for synchrony AT0 stimulation.

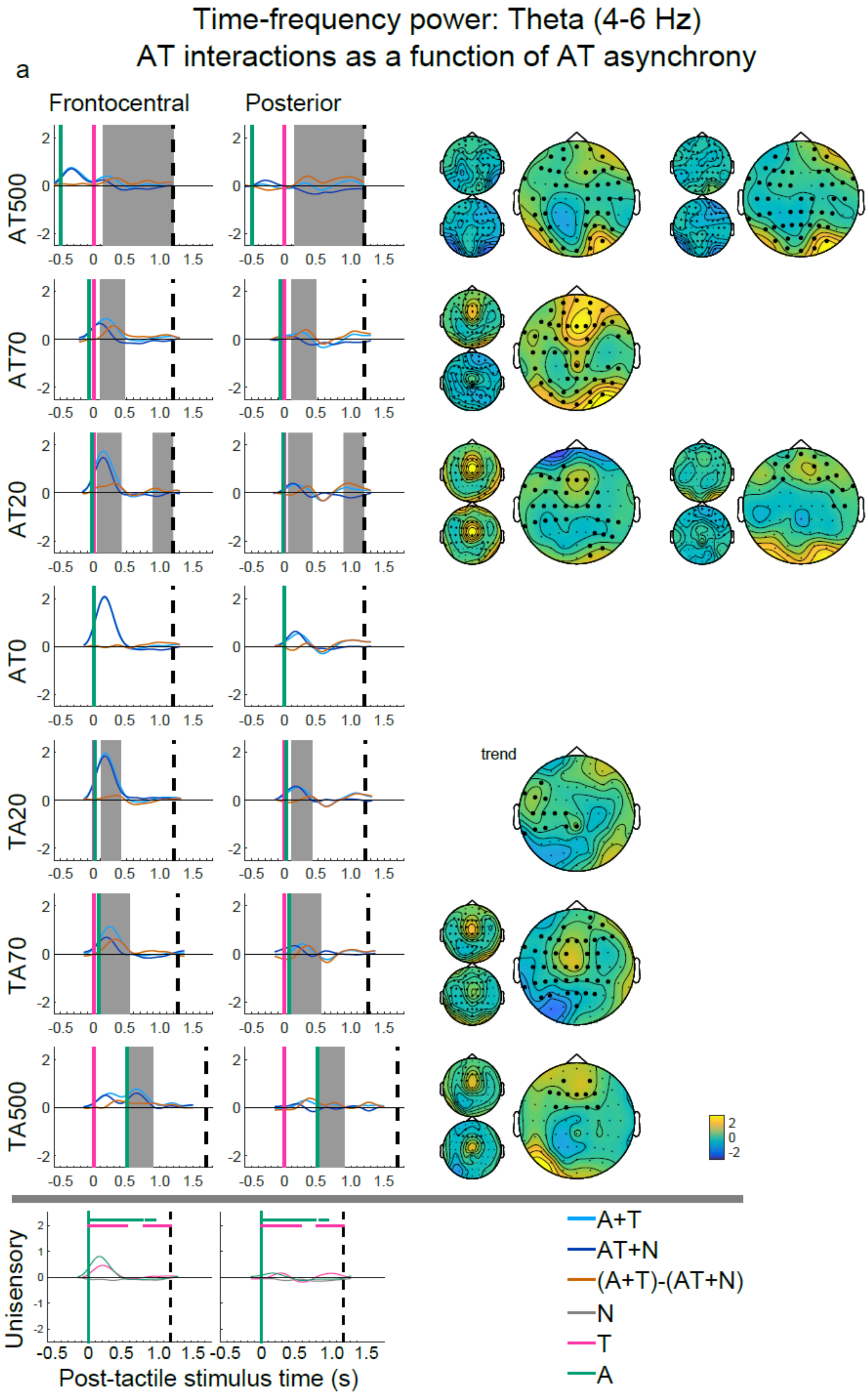

Time-frequency power: Theta (4-6 Hz)
AT interactions as a function of AT asynchrony
a
Frontocentral
Posterior
AT500
AT70
AT20
AT0
TA20
trend
TA70
TA500
Unisensory
Post-tactile stimulus time (s)
A+T
AT+N
(A+T)-(AT+N)
N
T
A

**Figure S7:** Theta-band power. Each row shows the audiotactile interaction for a particular level of AT asynchrony. (A) Theta power of the sum of the auditory and tactile (A+T; light blue), the sum of audiotactile plus null (AT+N; dark blue), and the audiotactile interaction, i.e. the difference ([A+T]-[AT+N], orange). The bottom row shows the null (grey), tactile (pink), and auditory (green) condition. Green = auditory onset, pink = tactile onset. Shaded grey areas indicate the timing of significant AT interactions at $p < 0.05$ corrected at the cluster level for multiple comparisons across electrodes and time points within a 1200 ms window starting with the second stimulus and limited by the black dashed line. Note that these effects were further corrected for 3 frequency bands (theta, alpha, beta) and are subject to a final threshold of $p<0.017$; any result with p-value between 0.017 to 0.05 is labelled as a 'trend'. (B) Topographies of the sums: A+T, AT+N, and (A+T)-(AT+N) for time windows of significant AT interactions, arranged in the same way as in Figures S2 and S3. A black star over an electrode indicates that it is part of a significant cluster. There are two separate columns for AT500 and AT20 as there were two distinct time windows of results for these two asynchronies.

In addition, we observed significant AT interactions for theta power in the AT500 and TA500 conditions. For AT500, in both early (60-600 ms) and late (610-1200 ms) time windows, AT interactions were found with topographies that were distinct from the fronto-central P200-like theta-band effects. For TA500, AT interactions were found only in the early (0-400 ms) time window after second stimulus, again with a topography distinct from the fronto-central P200 effect, yet similar to the early AT500 theta power effect.

*Beta power*: Both tactile and auditory stimuli induced changes in the beta band primarily over posterior channels (Figure S8, Unisensory row), which were more pronounced for tactile stimulation. Auditory and tactile stimulation initially suppressed beta power (event-related desynchronization; ERD) around 250 ms post-stimulation followed by a rebound (event-related synchronisation; ERS) above and beyond baseline, around 800-1000 ms post-stimulation; only the rebound was a significant change.

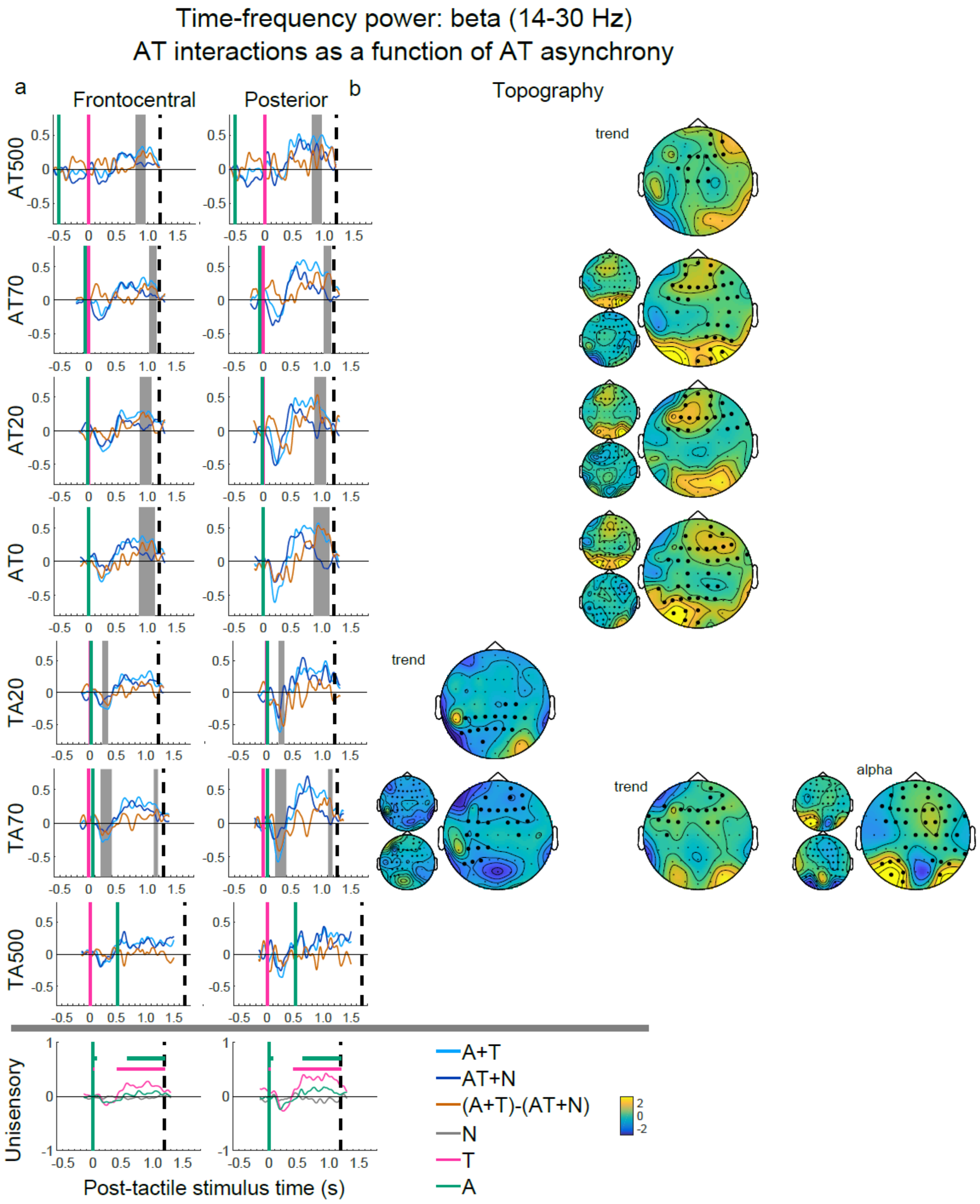


**Figure S8:** Beta-band power. Same details as for Figure S7. Topographies in sub-figure B are arranged in to 3 columns, namely the first to depict early (<500 ms) effects, the second for late (>500 ms) effects, and the third column for the one late alpha-band effect that was significant for TA70 and had a similar topography to the beta-band effect for TA70.

The early beta effect was altered by the AT+N condition relative to A+T, primarily for the TA70 condition, in which the A+T display a stronger suppression than AT+N, with a diffuse topography although strongest for posterior electrodes. This effect is further related to the rmANOVA results in Figure 5 and the main text.

The later beta power rebound was altered for AT + N relative to A + T across several asynchrony levels including AT70, AT20, AT0, and TA70 conditions (Figure S2 for statistics and Figure S8). Specifically, the rebound in alpha/beta power occurred earlier, was attenuated, and decayed faster for AT+N than the A+T sum, where alpha/beta power rebound was found to be more sustained (800-1100 ms post-stimulation).

Because the AT interactions of power rebound (~1000 ms) occurred after the explicit detection response is made by participants in the redundant target paradigm (~250 ms), it may be a consequence of the implicit AT event detection, or be associated with post-decisional processes such as metacognitive monitoring (Deroy et al. 2016), or the binding of asynchronous signals into a single multisensory percept (Roa Romero et al. 2015). Future redundant target paradigms that combine target detection with post-decisional tasks (e.g. confidence judgments) may enable us to further determine the functional role of the beta rebound and the associated AT interactions. The distinct response profile for theta versus beta power, varying with stimulus asynchrony, is in line with distinct mechanisms for different frequencies (Keil and Senkowski 2018).
